# Evaluation of Dose Fractionated Polymyxin B on Acute Kidney Injury: A Translational *In Vivo* Model

**DOI:** 10.1101/845024

**Authors:** Jiajun Liu, Gwendolyn M. Pais, Sean N. Avedissian, Annette Gilchrist, Andrew Lee, Nathaniel J. Rhodes, Alan R. Hauser, Marc H. Scheetz

**Affiliations:** Midwestern University, Downers Grove, IL; Midwestern University Chicago College of Pharmacy Pharmacometrics Center of Excellence, Downers Grove, IL; Northwestern Memorial Hospital, Chicago, IL; Antiviral Pharmacology Laboratory, University of Nebraska Medical Center (UNMC) Center for Drug Discovery, UNMC, Omaha, NE; University of Nebraska Medical Center, College of Pharmacy, Omaha, NE; Northwestern University, Chicago, IL

**Keywords:** polymyxins, polymyxin B, colistin, CMS, pharmacokinetic, toxicodynamic, animal model, acute kidney injury, biomarker, KIM1

## Abstract

The polymyxins are last-line defense for highly resistant infections. Nephrotoxicity, however, is a dose-limiting factor. Yet, approaches to mitigate nephrotoxicity are poorly defined. This study aimed to investigate the impact of dose fractionated (once, twice and thrice daily) polymyxin B (PB) on acute kidney injury (AKI) in a pre-clinical model. Secondarily, we aimed to describe the pharmacokinetic (PK) profile of PB. Sprague-Dawley rats were assigned to experimental groups with different dosing intervals but constant total daily exposure (12 mg/kg/day into single, twice daily, and thrice daily doses) and controls received normal saline subcutaneously over 3 days. Blood and urine samples were collected, and kidneys were harvested at necropsy. A three-compartment model best described the data and Bayesian observed vs. predicted concentration demonstrated bias, imprecision, and R^2^ of 0.129 mg/L, 0.729 mg^2^/L^2^ and 0.652, respectively. PB exposure (i.e. AUC_24h_) were similar across treatment groups over time (p=0.87). As a representative, urinary KIM-1 were elevated on days 1 and 2 for experimental groups compared to controls, and thrice daily group experienced the most KIM-1 increase [mean increase (95% CI) day 1 from day −1, 4.44 (0.89, 8.00) ng/mL; p=0.018] as compared to control [mean increase (95% CI) day 1 from day −1, 0.03 (−0.42, 0.49) ng/mL; p=0.99]. Correspondingly, significant histopathological damage was observed with the same group (p=0.013) (controls as a referent). Our findings suggested that fractionating the PB dose thrice daily resulted in the most injury in a rat model.

## Background

The widespread use of broad-spectrum antimicrobial agents has led to an increasing rate of resistant infections, and Gram-negative pathogens including *Pseudomonas aeruginosa, Acinetobacter* spp., and *Enterobacteriaceae* spp. are particularly problematic (1, 2). The Center for Disease Control and Prevention (CDC) has published a comprehensive report detailing the top antibiotic resistant threats in the U.S., stating that at least 2.86 million Americans contract resistant bacterial and fungal infections and at least 35,900 die annually as a result (3). It is estimated that by 2050, global deaths due to antimicrobial resistance will balloon to 10 million people per year and become the leading cause of mortality (4). Multiple Gram-negative species are resistant to nearly all available antibiotics (3), including newer combination agents (5, 6). Although multiple drugs in the antibiotic pipeline are promising, there is a prudent need to maximize clinical efficacy and safety of currently available agents (7). As a result of the paucity of active antibiotics for these difficult-to-treat infections, the polymyxins remain last resort options (8, 9).

The polymyxins are a group of polypeptide antibiotics discovered more than 70 years ago, with activity and efficacy against Gram-negative pathogens (10, 11). Polymyxin use has declined as a result of associated renal and neurological adverse effects and newer agents with more favorable safety profiles have emerged (12, 13). Thus, the treatment-limiting adverse effects of the polymyxins such as kidney injury have greatly limited their utility for patient care (14–16). Contemporary studies utilizing widely accepted dosing regimens have demonstrated that nephrotoxicity rates range from 21-48% (17–20). The mechanism for which nephrotoxicity develops appears to involve several processes. First, the drug is selectively reabsorbed by the renal brush border membrane and accumulates in renal cells, directly exerting cytotoxic effects to the proximal tubule cells (21). Secondly, accumulation of drug in the kidneys leads to increased membrane permeability and cell lysis, causing acute tubular necrosis (15, 22). Lastly, oxidative stress may also play a role in the development of nephrotoxicity associated with polymyxin therapy (16, 23, 24).

While the various mechanisms of polymyxin toxicity are being elucidated, dosing strategies that accelerate/diminish toxicity remain poorly defined. Toxicity thresholds for plasma 24-hour area under the curve (AUC_24h_) of polymyxin B and colistin have recently been highlighted, yet approaches to minimize nephrotoxicity risk resulted in mixed outcomes (20, 25–29). More specifically, it remains unclear whether dividing the total daily dose of polymyxins into fractions (e.g. giving twice or thrice daily) can circumvent kidney injury during treatment. In this study, we examined the impact of dose fractionated systemic polymyxin B on acute kidney injury (AKI) in a pre-clinical, humanized model with novel urinary biomarkers (30, 31). In addition, we aimed to describe the polymyxin B pharmacokinetic (PK) profile.

## Material and Methods

### Chemical and reagents

Clinical grade Polymyxin B sulfate (PB) (Lot# CD807) for injection (USP) was purchased from X-GEN Pharmaceuticals (Horseheads, NY, USA). Study drug was reconstituted and diluted with normal saline (NS) for injection. Unused portions were properly discarded to minimize loss of potency in subsequent experiments (32). Colistin sulfate (Sigma-Aldrich Chemical Company, Milwaukee, WI, USA) and creatinine-d3 (Cayman Chemicals, Ann Arbor, MI, USA) were used as internal standards for the LC-MS/MS assay (Agilent Technologies). All solvents for the LC-MS/MS analysis were of LC-MS/MS grade. Acetonitrile and methanol were purchased from VWR International (Radnor, PA, USA). Formic acid was obtained from Fisher Scientific (Waltham, MA, USA). Pooled male Sprague-Dawley rat plasma was used for sample preparation and calibration of standard curves (BioreclamationIVT, Westbury, NY, USA).

### Experimental design and animals

Allocation and number of animals for experimental (i.e. PB-treated) and control protocol groups are described in Fig. 1. The experimental arm was further divided into three groups based on dose fractionation design: once daily (QD), twice daily (BID), and thrice daily (TID). Each experimental group received subcutaneous injections of PB, and control groups received equal volumes of NS based on the QD protocol. In all, there were four study groups. Total daily dose of PB was fixed at 12 mg/kg/day (allometrically scaled) for all experimental groups administered subcutaneously for 72 hours (i.e. 3 PB doses for QD group, 6 for BID group, and 9 for TID group during the 3-day study period) (33).

**Figure 1.**
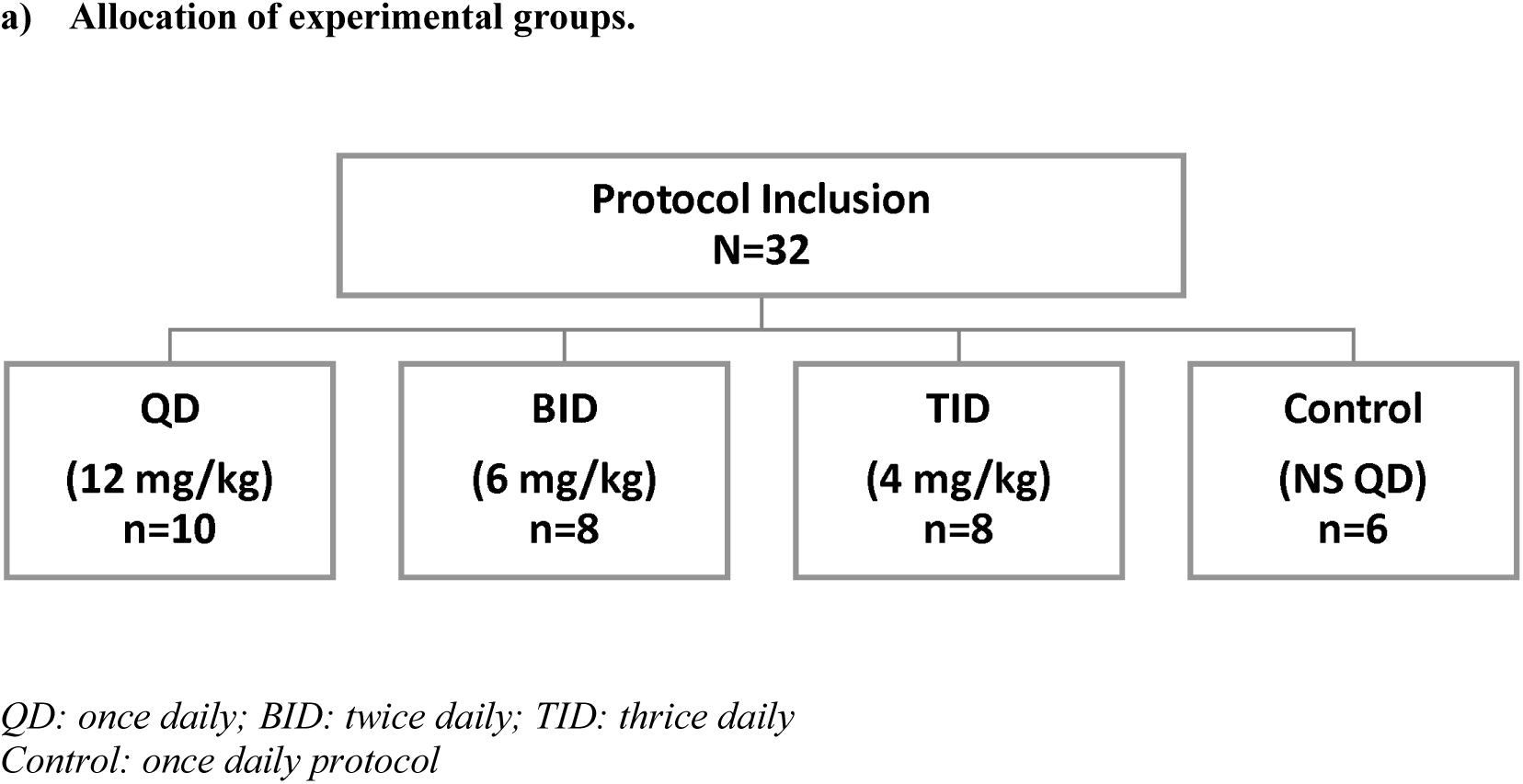

Male Sprague-Dawley rats (n=32, approximately 8 to 10 weeks old; Harlan, Indianapolis, IN, USA) were used. Animals were housed in a light- and temperature-controlled rooms during acclimation and study periods. Animals were maintained in plastic cages on a 12-hour light and 12-hour dark cycle. Food and water were freely accessible at all times except during periods in which sampling catheters (one per animal) were surgically placed prior to initiation of study protocol. Post-operative pain was monitored according to protocol. Data were analyzed for all protocol-initiated animals unless terminated early, in which case data were treated as missing.

This study was conducted at Midwestern University, Downers Grove, IL. The study methods were reviewed and approved by the Midwestern University Institutional Animal Care and Use Committee (IACUC; protocol 2677). All animals were cared for and handled in concordance with animal care and use standards and ethical principles.

### Blood and urine sampling

Fig. 2 provides a schematic flow of study design for each study group. Blood samples were obtained via an internal jugular vein catheter that was surgically cannulated on day 0 after the acclimation period. Animals were under ketamine (100 mg/kg) and xylazine (10 mg/kg) anesthesia for surgical procedure and were allowed 24-hour recovery periods prior to initiation of the study protocol (i.e. drug dosing or blood sampling). When not in use, catheters were locked with heparin solution (100 IU/mL). Blood samples (0.125 mL aliquots) were obtained after the first dose (day 1) with a staggered sampling design. A maximum of 16 samples per animal were obtained during a 4-day period (pre-euthanasia) and no more than 8 samples were drawn in a single day. As an example, for QD and control groups blood samples were drawn at 5, 20, 60, 120, 180, 240, 360, and 480 minutes after the first dose on day 1. Eight blood samples total were then obtained on days 2 and 3. A terminal sample was also drawn under terminal anesthesia. Blood sampling schemes for BID and TID groups follow the same protocol (i.e. maximum number of samples and volume per animal) while the sampling times were adjusted accordingly based on the dosing intervals. Each sample was replaced with equal volume of NS to maintain euvolemia. Blood samples were immediately transferred to a disodium EDTA (Sigma-Aldrich Chemical Company, Milwaukee, WI, USA) treated microcentrifuge tube and centrifuged at 600 *xg* for 10 minutes. Plasma supernatant was collected and stored at - 80°C for batch sample analysis. Animals were placed in metabolic cages for urine collection as previously described (34, 35). In brief, discrete entry times were recorded for initial transfer of animals to the metabolic cages (catalogue number 650-0350; Nalgene, Rochester, NY) on day −1 (baseline), and urine collections and volume measurements followed the 24-hour period on days 2, 3 and 4. All urine samples were collected in laboratory-controlled ambient conditions, and urinary biomarkers were stable throughout as previously described (35). Urine samples were centrifuged at 500 *xg* at 4°C for 10 minutes, and supernatant was stored at −80°C for batch analysis.

**Figure 2.**
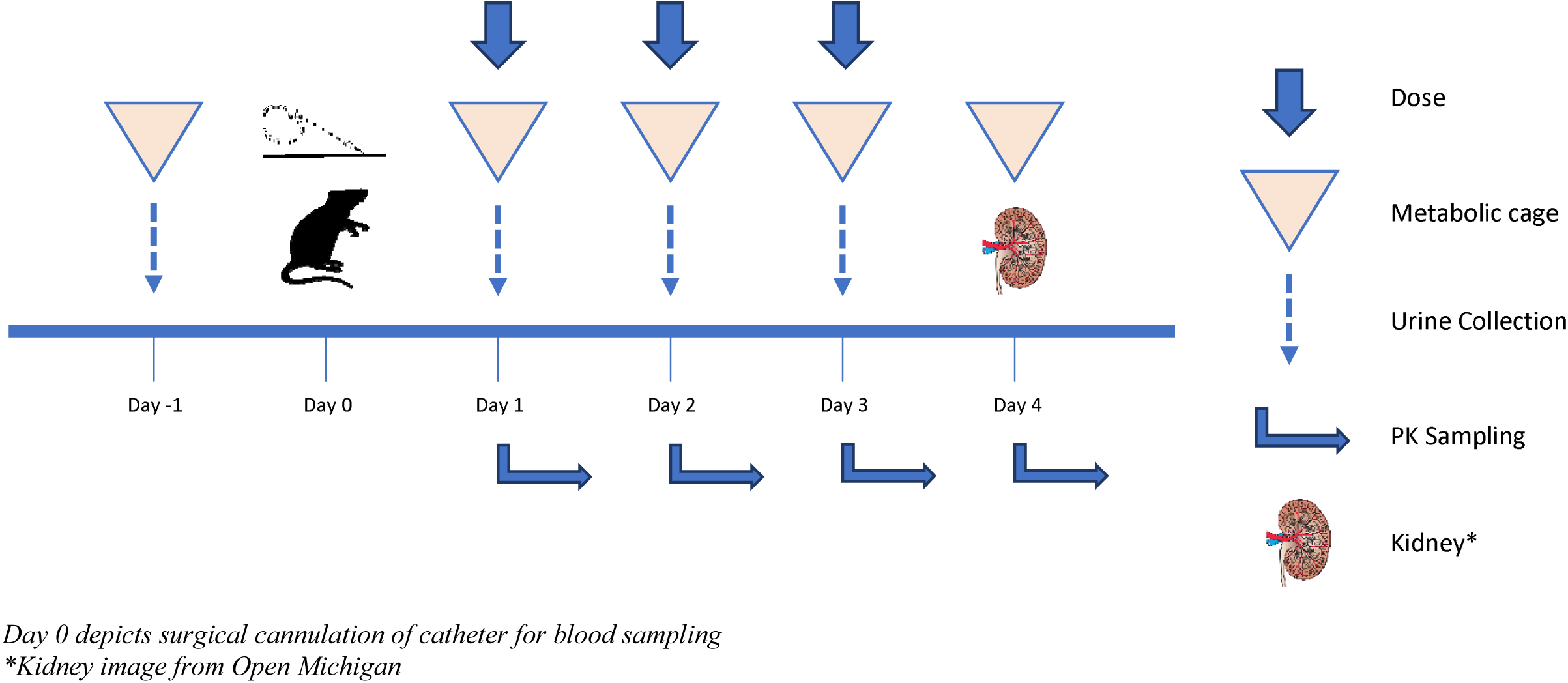
Schematic of study design for each study group.

### Determination of PB and creatinine concentrations in plasma

For quantification of plasma PB concentration, 40 μL plasma sample was combined with 4 μL of internal standard of colistin sulfate at a concentration of 0.1 mg/mL. Protein precipitation was then performed with 456 μL of methanol containing 0.1% formic acid. Following centrifugation for 10 minutes at 16,000 *xg* (Eppendorf model: 5424), 100 μL supernatant was collected for analysis. Processed samples were injected (injection volume at 2 μL) into an Agilent 1260 infinity binary liquid chromatograph paired with Agilent 6420 triple quadrupole mass spectrometer (MS). A Poroshell 120 EC-C18 column (100 mm x 3 mm, 2.7 μm) was used. The following quantifier transitions (m/z) for polymyxin B1 (PB1) and colistin A were identified and utilized: 402.2 ➔ 101.1, 390.6 ➔ 101.3, respectively. The assay was linear between 0.5 to 40 mg/L (R^2^=0.997) for PB1 after an applied weight of 1/x. To quantify plasma creatinine, m/z transitions of 117.09 ➔ 89.2, 114.1 ➔ 44.3 were utilized for creatinine-d3 and creatinine, respectively. After adjusting for endogenous creatinine in pooled blank plasma and 1/x weighting, the linear range for creatinine assay was between 0.3 and 40 mg/dL (R^2^=0.999). The coefficient of variation (CV%) values for PB1 and creatinine assays were below 10% for intra- and inter-day measures. All samples measuring above the upper limit of quantification underwent serial dilution for analysis.

### Determination of urinary biomarkers of kidney injury

Urine samples were analyzed for creatinine content and urinary biomarkers. Urine aliquots were analyzed in batches to determine the concentration of KIM-1, IP-10, TIMP-1, CLN and OPN. Urinary biomarkers were assayed using a microsphere based MAGPIX kit as previously described (35). In brief, urine samples were aliquoted into 96-well black plates supplied with MILLIPLEX^®^ MAP Rat Kidney Toxicity Magnetic Beed Panel 1 (EMD Millipore Corporation, Charles, MO, USA), prepared and analyzed per manufacturer’s recommendations.

### Histopathological examination of renal cell damage

Kidney tissues were harvested following euthanasia. Each animal’s kidneys were removed and briefly washed in cold NS. The left kidney was fixed in 10% formalin for histopathology examination. Histopathological analysis of kidneys (n=32) was conducted by IDEXX BioAnalytics (Columbia, MO, USA). A validated, ordinal scoring system was employed to grade pathological lesions as previously described (31, 36, 37). Briefly, a scale of 0 indicates no abnormality while a scale of 5 indicates massive and extensive renal damage. The final histopathology score for an individual animal was calculated based on the highest score from the anatomical structural segment.

### PB pharmacokinetic model and exposure determination

To construct the base PK models and generate exposure estimates for each individual animal, the Nonparametric Adaptive Grid (NPAG) algorithm (38, 39) within the Pmetrics (Version 1.5.2) package (39) for R (6) was utilized. Multiple models were built and assessed. A three-compartmental structural model of PB disposition accounting for absorption constant (K_a_) from injection site to the central compartment (V_C_) was fitted to all PK data. The PK model was parameterized with K_a_, V_C_, intercompartmental transfer rates (K_23_, K_32_) between central and peripheral (V_P_) compartments, and total elimination rate constant (K_e_). Assay error was included in the model using a polynomial equation in the form of standard deviation (SD) as a function of each observed concentration, Y (i.e. SD = C0 + C1 * Y). Observation weighting was performed using lambda (i.e. error = SD^2^ * lambda^2^)^0.5^, an additive variance model to account for extra process noise. Lambda was initially set at 1 with C0 and C1 equal to 0.1 and 0.1, respectively. Comparative model performance was examined by the change in objective function value (OFV) calculated as differences in −2 log-likelihood (−2LL), with a reduction of 3.84 in OFV corresponding to p <0.05 based on chi-square distribution with one degree of freedom. Further, the best-fit model was selected based on the rule of parsimony and the lowest Akaike’s information criterion (AIC) scores. Goodness-of-fit of the competing models were evaluated by regression on observed vs. predicted plots, coefficients of determination, and visual plots of individual Bayesian predicted concentration-time profiles. Bias was defined as mean weighted prediction error; imprecision was defined as bias-adjusted mean weighted squared prediction error. Using the final model, PB exposure indices (AUC_24h_, C_MAX_, and C_MIN_) were calculated from individual Bayesian posterior-predicted concentrations using ‘makeNCA’ within Pmetrics package across 24-hour intervals (39).

### Statistical analysis for biomarkers, PK indices and histopathological scoring

Analysis of variance (ANOVA) with Geisser and Greenhouse epsilon hat correction method accounting for subjects, treatment groups, repeated measures over time or a mixed-effects model (when data were missing) were utilized for statistical analyses of urinary biomarkers, plasma creatinine, and PK indices between study groups using GraphPad Prism (version 8.2.1 for Windows, GraphPad Software, La Jolla, CA). Ordinal logistical regressions on histopathological scores were performed with observed nominal scores treated as dependent variables and control group as the referent category using Stata version 13 (40). All tests were two-tailed with an α level of 0.05 for statistical significance.

## Results

### Differences between animal cohorts

A total of 32 PB-treated animals contributed PK model data and completed all protocols. All animals weighed between 291.1 to 321.1 g. Pre-surgery (i.e. day −1) mean (SD) urine volumes were 5.2 (1.7), 8.6 (2.2), 6.4 (2.3), and 5.8 (2.2) mL for QD, BID, TID and control groups, respectively. The QD group had significantly lower urine volume compared to BID group at day −1 (p=0.01), while no statistical differences in urine output were observed between experimental groups on days 1, 2, and 3 (Fig. 3). All experimental groups produced significantly more urine compared to controls on study days 1, 2 and 3, except BID group on day 1 (p=0.23).

**Figure 3.**
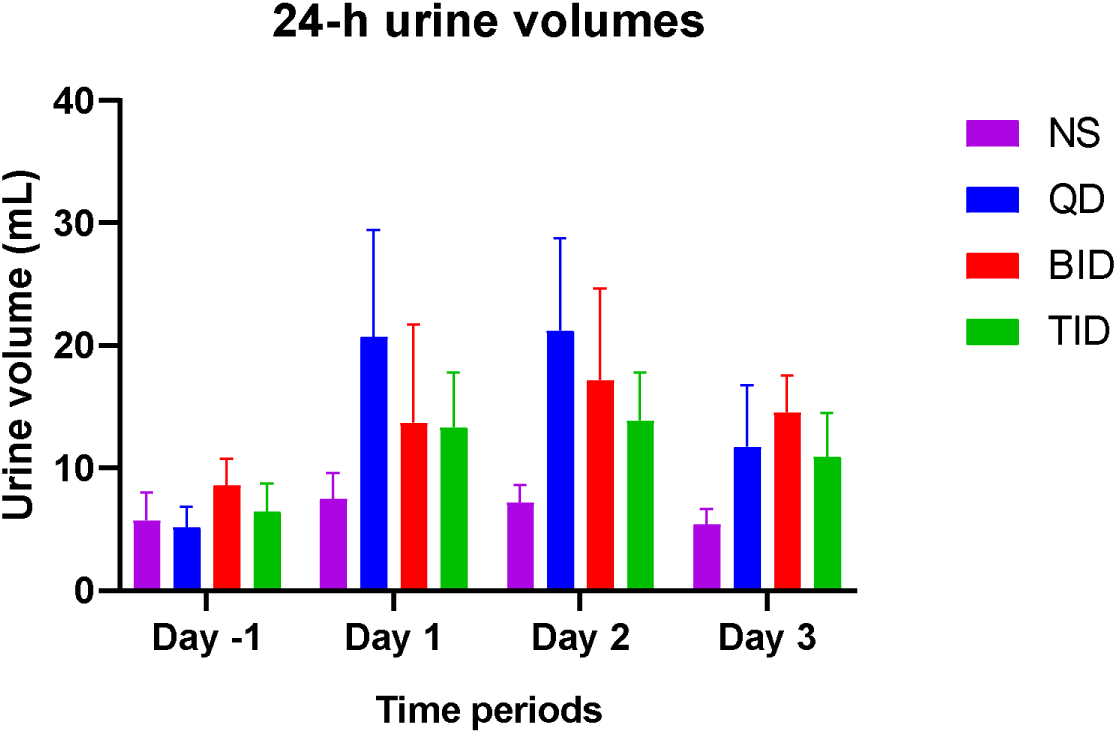
Mean 24-h urine volume between experimental and control groups.

### PB pharmacokinetic models and exposures

Various modeling approaches were utilized to fit the PK data. A three-compartment model was chosen as the final model given that it was the most parsimonious with the least bias and imprecision and displayed the most significant OFV change with the lowest AIC against competing models (Table 1). Table 2 provides a summary of the population mean parameter values for K_a_, V_C_, K_23_, and K_32_. Model predictive performance for observed vs. Bayesian posterior-predicted concentrations for bias, imprecision, and R^2^ were: 0.129 mg/L, 0.729 mg^2^/L^2^ and 0.652, respectively (Fig. 4). PK exposures were calculated based on NCA analysis on the Bayesian posterior-predicted concentrations from the best-fit PK model and are graphically represented in Fig. 5. Mean AUC_24h_ (SD) for QD, BID, and TID groups were 171.1 (41.0), 168.6 (58.9), and 129.6 (50.7) mg*h/L, respectively. Similarly, mean (SD) C_MAX_ for QD, BID, and TID groups were 10.7 (0.80), 9.7 (0.47), and 6.7 (0.49) mg/L, respectively and mean C_MIN_ (SD) were 2.3 (1.3), 3.7 (1.4), and 3.2 (1.2) mg/L, respectively. Statistical procedures were conducted to evaluate the differences in AUC_24h_, C_MAX_ and C_MIN_ between experimental groups and respective 24-hour intervals. Although the QD group exhibited higher AUC_24h_ than TID group (p=0.003) on day 1, no significant effects were observed for overall exposure vs. time (p=0.87). Similarly, the QD group had an overall lower mean C_MIN_ compared to the BID group (p <0.0001) or TID group (p<0.0001) during the first 24 hours, but no significant differences were observed between experimental groups over the entire study period (p=1.00). Compared to the TID group, both QD and BID groups showed significantly higher C_MAX_ during the first 24 hours (p=0.0083 and p=0.049, respectively). The QD group exhibited the highest C_MAX_ on days 2 and 3 (p=0.0025 and p=0.0017, respectively). On days 2 and 3, the C_MAX_ in the BID group was numerically elevated when compared to the TID group, but the values were not statistically significant (p= 0.051 and p=0.055, respectively).

**Figure 4.**
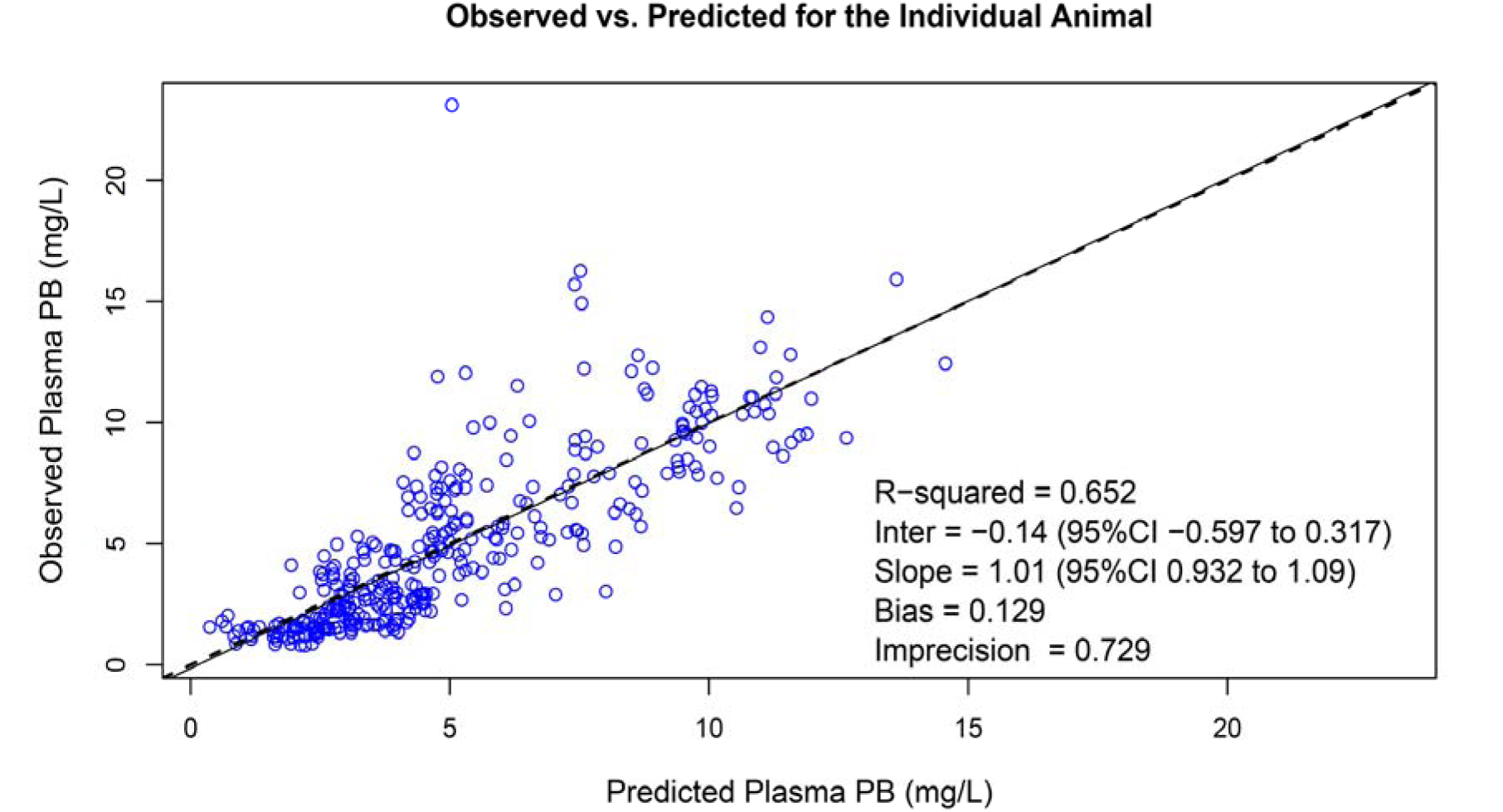
Goodness-of-fit plot for Bayesian observed vs. predicted plasma PB concentrations utilizing the final three-compartment model.

**Figure 5.**
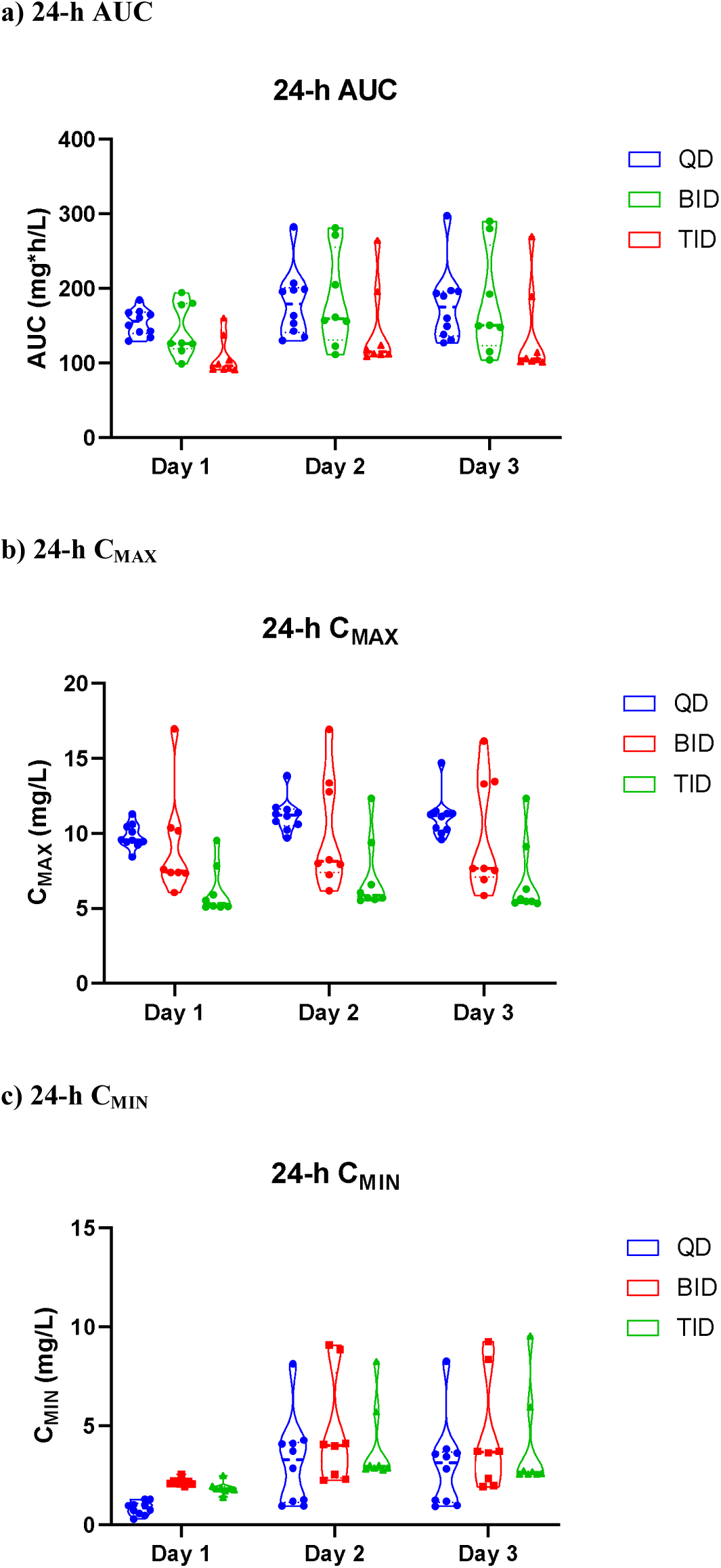
Violin plots of PK indices from the final best-fit model by days.

**Table 1.** Model selection summary.

| **Compartmental Models** | **-2LL** | **OFV Change** | **AIC** | **Bias (mg/L)** | **Imprecision (mg<sup>2</sup>/L<sup>2</sup>)** | **Bayesian R<sup>2</sup>** |
| --- | --- | --- | --- | --- | --- | --- |
| **One** | 1709 | Ref | 1715 | -0.48 | 0.95 | 0.10 |
| **Two** | 1709 | 0 | 1720 | -0.05 | 0.90 | 0.20 |
| **Three** | 1450 | 259 | 1463 | 0.13 | 0.73 | 0.65 |

**Table 2.** Mean population parameters from the final model.

|  | Mean | SD |
| --- | --- | --- |
| $K_a$ (h<sup>-1</sup>) | 0.290 | 0.460 |
| $K_e$ (h<sup>-1</sup>) | 0.411 | 0.076 |
| $V_C$ (L) | 0.056 | 0.079 |
| $K_{23}$ (h<sup>-1</sup>) | 6.214 | 11.430 |
| $K_{32}$ (h<sup>-1</sup>) | 3.163 | 43.00 |

### Kidney injury biomarkers and histopathological examinations

Plasma creatinine and urinary biomarkers are graphically represented in Fig. 6. Plasma creatinine did not differ across treatments over time (p=0.18); however, the following biomarkers showed a significant treatment effect over the study period: KIM-1 (p<0.0001), OPN (p=0.029), IP-10 (p=0.046), and TIMP-1 (p<0.0001). As a representative biomarker, KIM-1 rose rapidly on days 1 and 2 for all experimental groups. Notably, the TID group experienced the largest KIM-1 increase [mean increase (95% CI) day 1 from day −1, 4.44 (0.89, 8.00) ng/mL; p=0.018] as compared to control [mean increase (95% CI) day 1 from day −1, 0.03 (−0.42, 0.49) ng/mL; p=0.99]. Increases were also observed with QD [mean increase (95% CI) day 1 from day −1, 2.58 (−0.12, 5.27) ng/mL; p=0.06] and BID groups [mean difference (95% CI) day 1 from day −1, 0.84 (−0.05, 1.73) ng/mL; p=0.06]. Further, the TID group exhibited a significant KIM-1 increase on day 2 [mean increase (95% CI), 2.44 (1.22, 3.67) ng/mL; p=0.0013] and a nonsignificant decrease was observed on day 3 [mean decrease (95% CI), 2.39 (−0.053, 4.83) ng/mL; p=0.055). Mean KIM-1 changes did not differ on days 2 and 3 between QD and BID groups. Similar trends and significant treatment and time effects were also observed for OPN (p=0.029), IP-10 (p=0.046), and TIMP-1 (p<0.0001) but no significant effects were observed with CLN (p=0.093) (Fig. 6).

**Figure 6.**
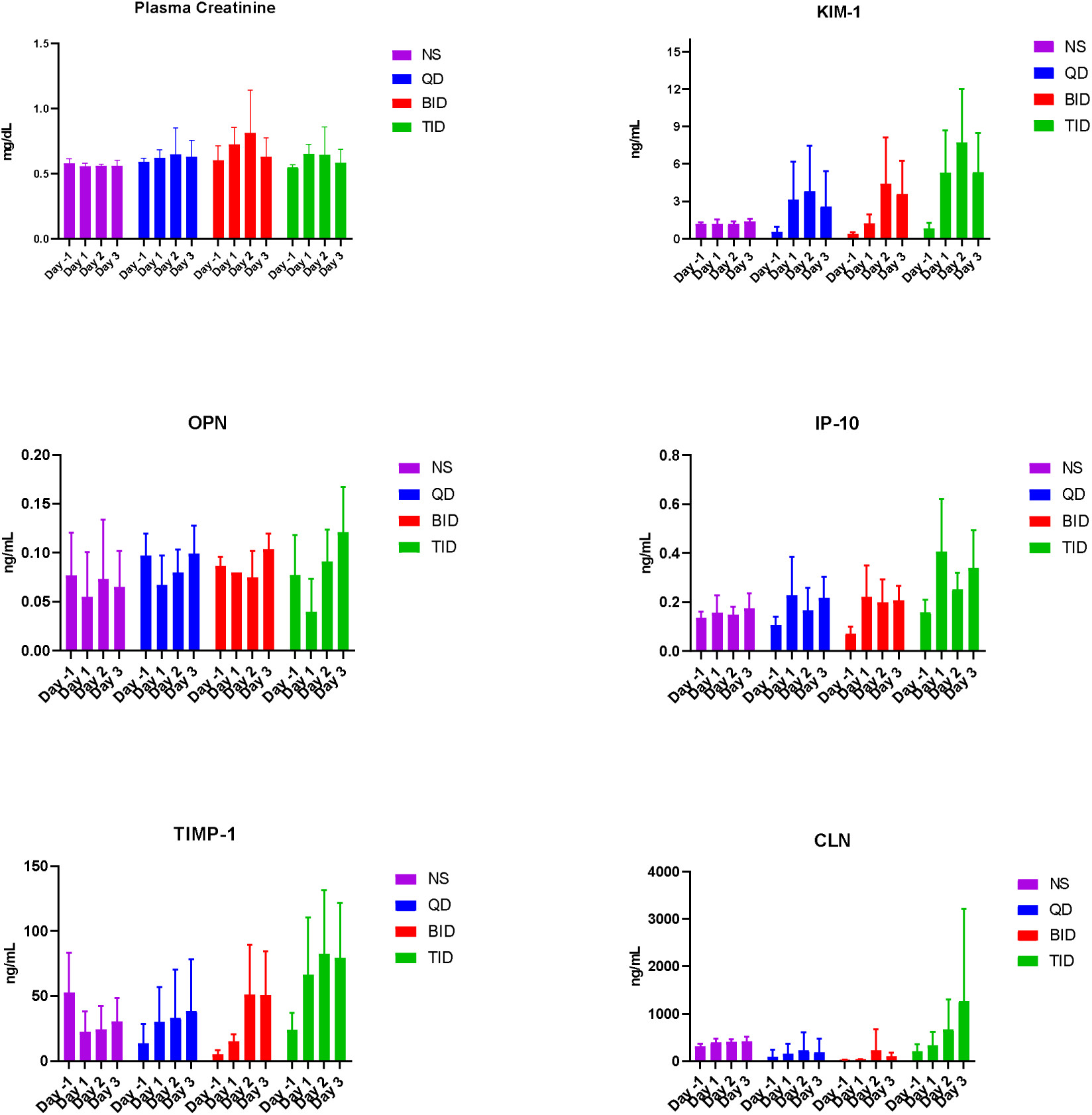
Plasma creatinine and urinary biomarkers.

Histopathological scorings are summarized in Table 3 and graphically displayed in Supplemental Figure S1. Representative histopathology images are provided in supplemental materials (Figures S2, S3, S4 and S5). Significant histopathological damage was observed with the TID group with a median [range] score of 2 [2, 3.5] (p=0.013; 95% CI, 0.70, 5.88) using controls as a referent category. While damage was also observed with QD [median score (range), 2 (1, 3)] and BID [2 (1, 3)] groups, no statistically significant difference was found when comparing either group to the referent group [p=0.156 (95% CI, −0.61, 3.77), p=0.092 (95% CI, −0.34, 4.50), respectively].

**Table 3.** Ordinal logistic regression of histopathology scorings by groups.

|  | Median Score [range] | p value | 95% CI |
| --- | --- | --- | --- |
| Control | 1.5 [1-2] | Referent | Referent |
| QD | 2 [1-3] | 0.156 | -0.61-3.77 |
| BID | 2 [1-3] | 0.092 | -0.34-4.50 |
| TID | 2 [2-3.5] | 0.013 | 0.70-5.88 |

## Discussion

A dose fractionation scheme in this study effectively maintained constant exposure and separated maximal concentrations across experimental groups. Our data demonstrated that fractionating the PB dose into three daily aliquots resulted in the most kidney injury as measured by the urinary biomarkers: KIM-1, OPN, IP-10, and TIMP-1. Significant differences were detectable within 24 hours. These markers have previously demonstrated high sensitivity for kidney injury (41–43). Further, KIM-1 and OPN were directly linked to proximal tubular toxicity, and this is concordant with previous reports (25, 41). Histopathological findings also indicated that thrice daily dosing of PB led to the most severe kidney insults within the study period. While the differences in urinary biomarkers and categorical damage scales were not significant for QD and BID groups, both groups demonstrated less extensive kidney injuries consistent with results from urinary biomarkers such as elevations in KIM-1 levels. Additionally, we derived a best-fit PK model for PB in rats using rich PK data. Utilizing the best-fit model, we found that the PB exposure (i.e. AUC_24h_) derived were similar across all experimental groups (QD, BID, and TID) and separation of peak concentrations was observed.

Contemporary dosing of intravenous PB recommends it be administered in 2 divided doses based on a weight-based total daily dose (26, 44). It has been suggested that PB-induced kidney injury could be minimized by optimizing dosing intervals, similar to that observed with aminoglycosides (7, 45). Wallace *et al*. utilized a preclinical rat model to explore this possibility for colistin. The authors found that the colistin methanesulfonate (CMS, prodrug of colistin) regimen corresponding to once daily dosing in humans led to a greater number and severity of renal lesions when compared to the group received fractionated dosage corresponding to twice daily dosing in human. They concluded that extended interval dosing of CMS resulted in more extensive renal damage (27). Abdelraouf *et al*. also utilized a rat model and administered PB subcutaneously at 20 mg/kg/day or 5 mg/kg every 6 hours (25). In contrast, they found a lower rate of nephrotoxicity associated with PB in the once daily group, while the split-dosage group experienced a quicker onset of nephrotoxicity (defined by elevation of creatinine from baseline); however, the rate of nephrotoxicity converged between groups towards the end of study. The authors suggested that an increased active, saturable, carrier-mediated uptake may have been responsible for this effect when PB was given repeatedly (as opposed to once daily) and led to the higher rate of renal injury. Most recently, Okoduwa *et al*. conducted a retrospective, propensity-score matched clinical study on 200 patients who received once-daily or twice-daily systemic PB across different medical centers (28). In contrast to the animal study by Abdelraouf *et al*., they found that a higher proportion of nephrotoxicity (using clinical criteria and considering all stages of AKI) was observed in the once-daily group than the twice-daily group (47% vs. 17%, respectively; p=0.0005). The findings from Okoduwa *et al*. are consistent with the animal model employed by Wallace *et al*., though CMS (and not PB) was utilized; however, it is unclear whether patients receiving once daily PB in clinical studies actually received equivalent (or greater) exposures compared to those receiving twice daily dosing. Our findings agree with the results from Abdelraouf *et al*. that dose fractionated PB strategy led to more extensive AKI. Additionally, our study provided rich PK data to confirm the exposure status across experimental groups and employed highly sensitive and specific urinary biomarkers for early detection of AKI in addition to plasma creatinine (46). The level of injury was further confirmed by histopathological examination.

We acknowledge several limitations to our study. First, our study was limited to 72-hour dosing compared to the relatively longer study period (up to 10 days) by Abdelraouf *et al*. The shorter time frame did not allow us to observe levels of plasma creatinine nor urinary biomarkers beyond this time frame; however, the urinary biomarkers utilized are highly sensitive in detecting early stages of AKI and our histopathology examinations confirmed the injury (30, 31, 46–48). Secondly, PB exposure (i.e. AUC_24h_) was held constant in our study and linking exposure to toxicodynamic data (i.e. injury biomarkers) would not be ideal as this was not an objective of our study. Thus, further studies are warranted to examine the PK/PD indices to toxicodynamic outcomes. Thirdly, this is a pre-clinical model and additional translational studies defining the lower limit of PB toxicity are needed to design maximally safe and effective dosing regimens.

To date, this is the first study that employed a rat model with a rich PK sampling design with dose fractionated systemic PB that also allowed PK estimates at an individual level. We also demonstrated that TID dosing of PB induces AKI as early as 24 hours. These findings may have clinical implications for PB dosing schemes in difficult-to-treat infections while minimizing nephrotoxicity. Further studies are warranted to explore PB exposure linked to toxicity while maximizing efficacy.

## Acknowledgements

None

## Supplemental Materials

**Figure S1.**
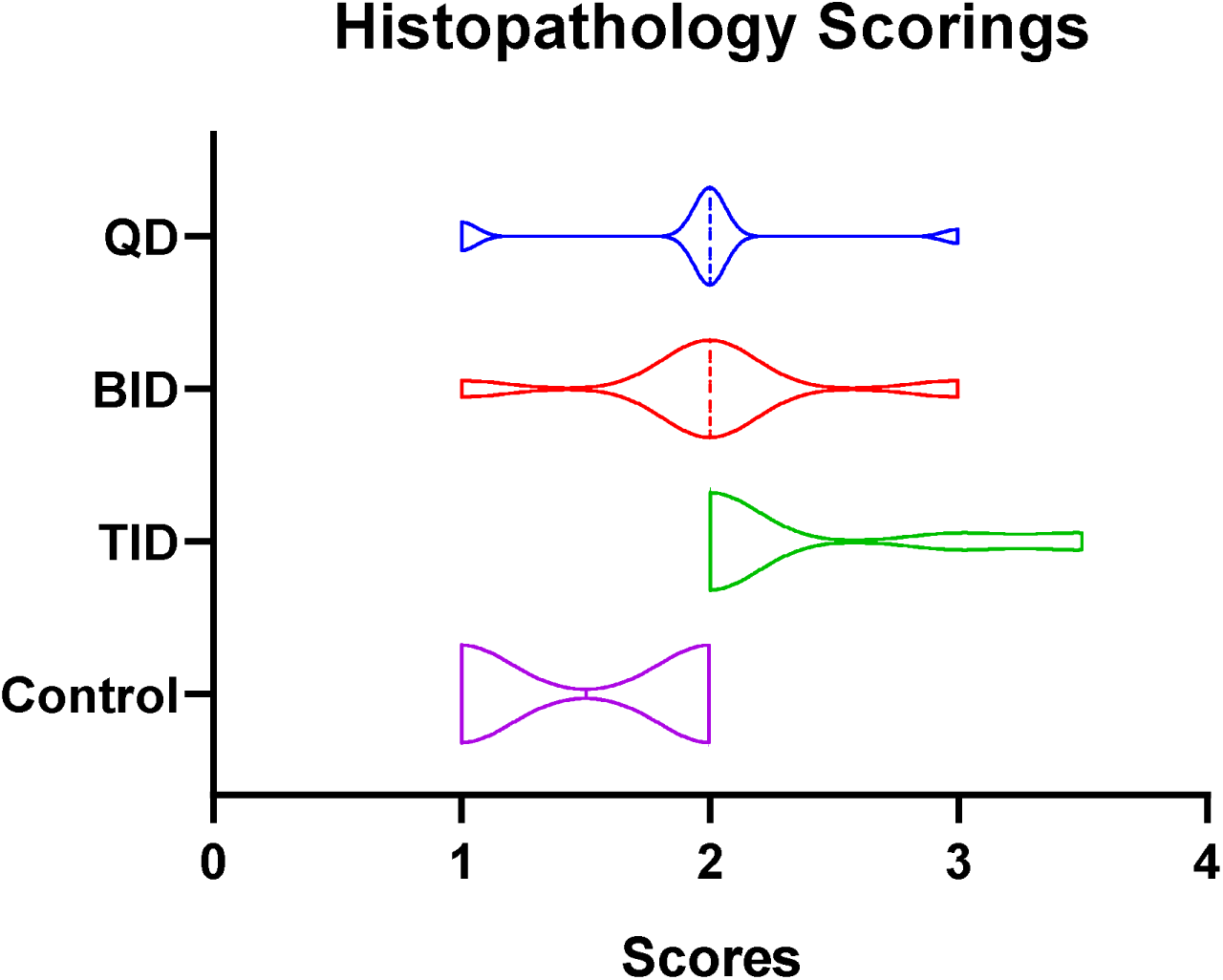
Violin plot of histopathological scores between all groups.

**Figure S2.**
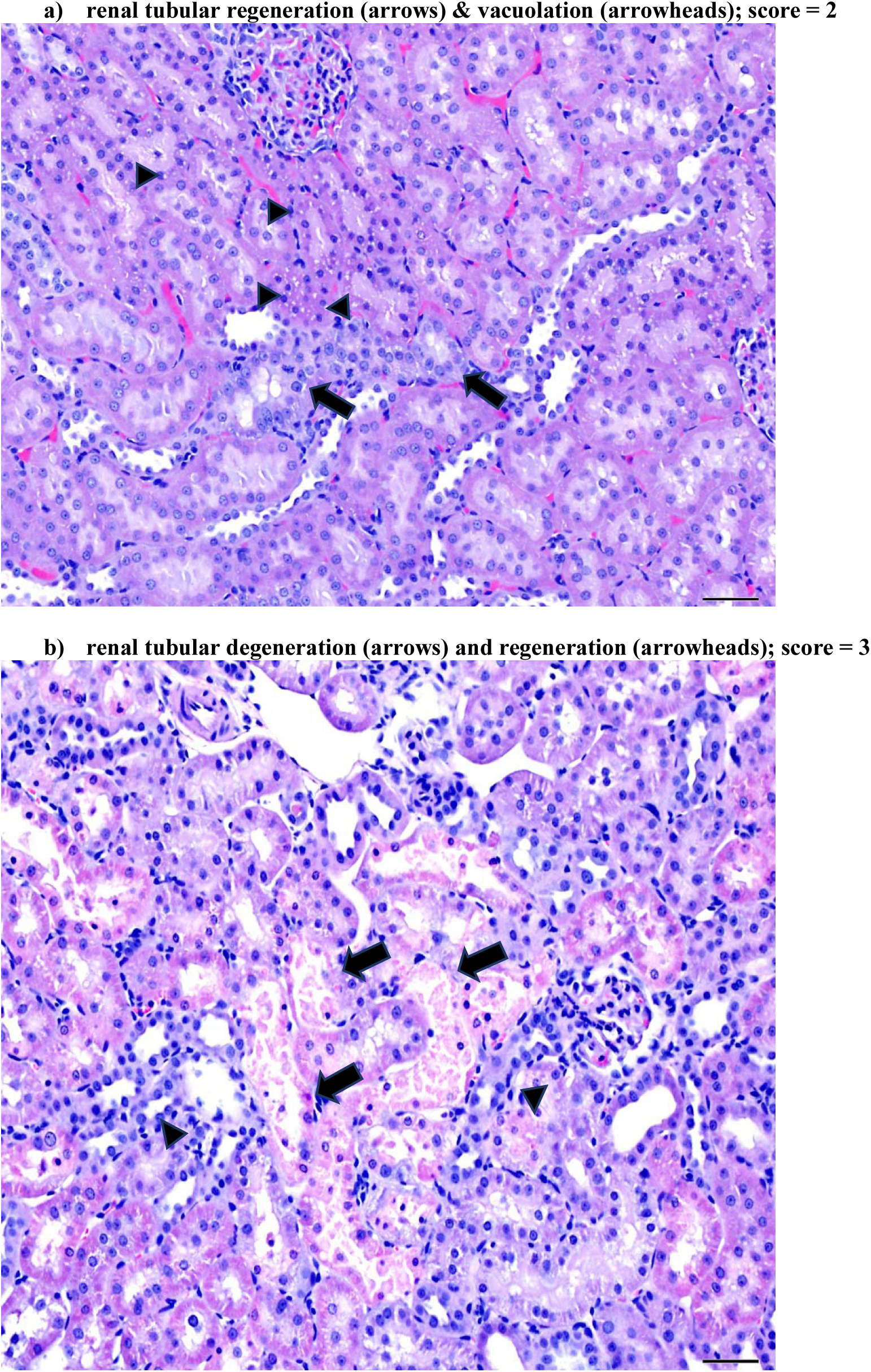
QD group histopathological examination images.

**Figure S3.**
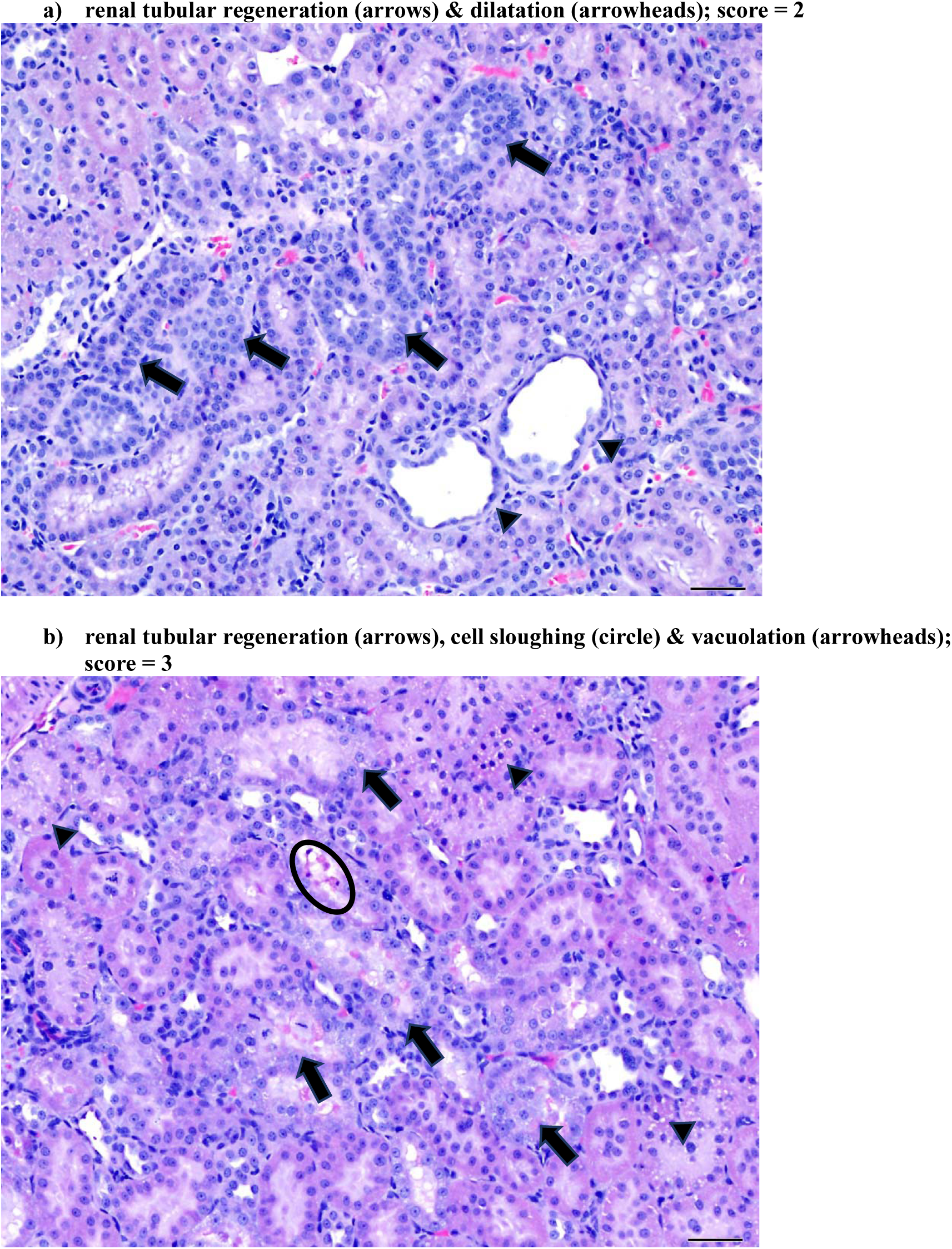
BID group histopathological examination images.

**Figure S4.**
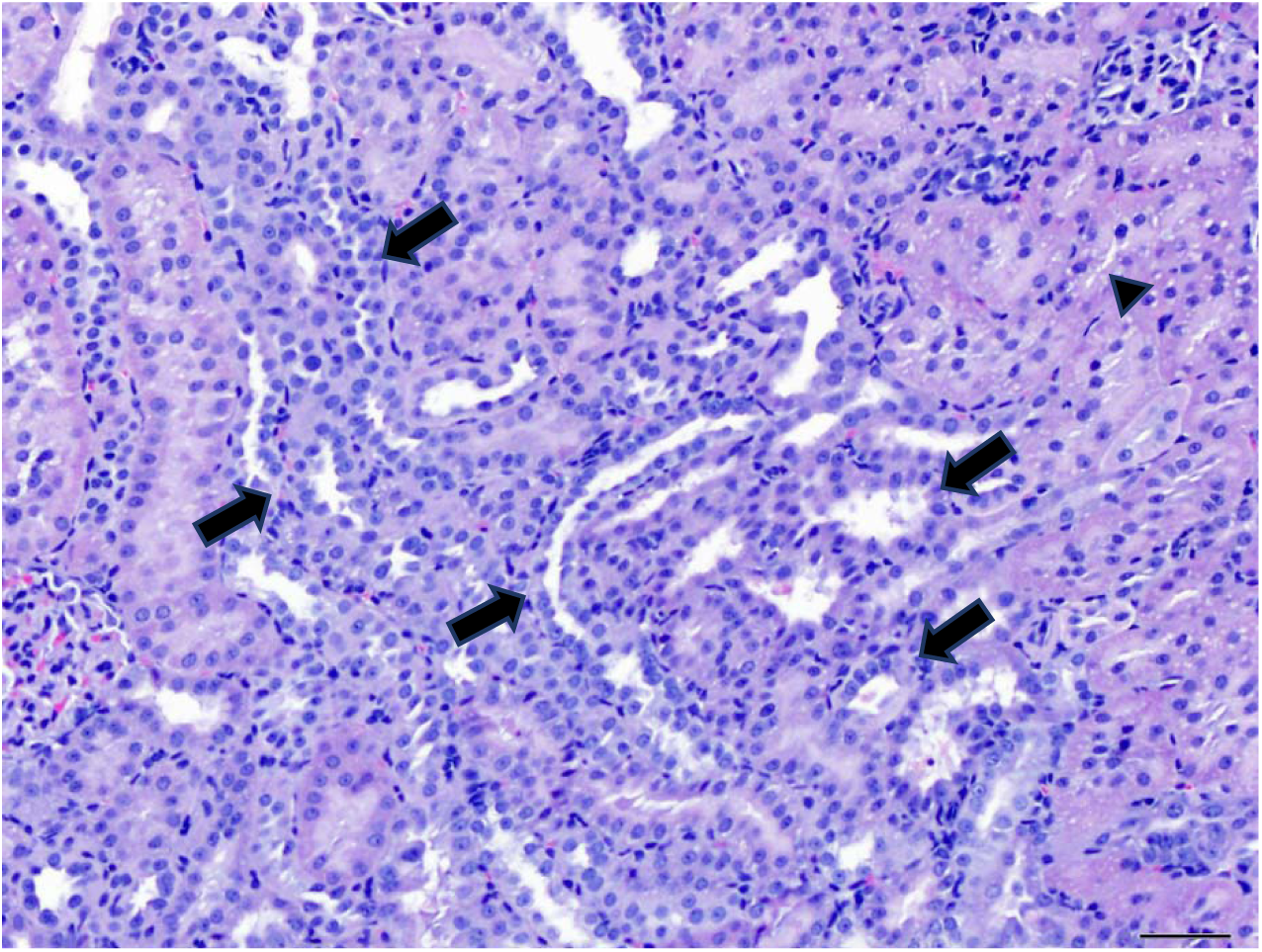
TID group histopathological examination images: renal tubular regeneration (arrows) & vacuolation (arrowheads); score = 3.

**Figure S5.**
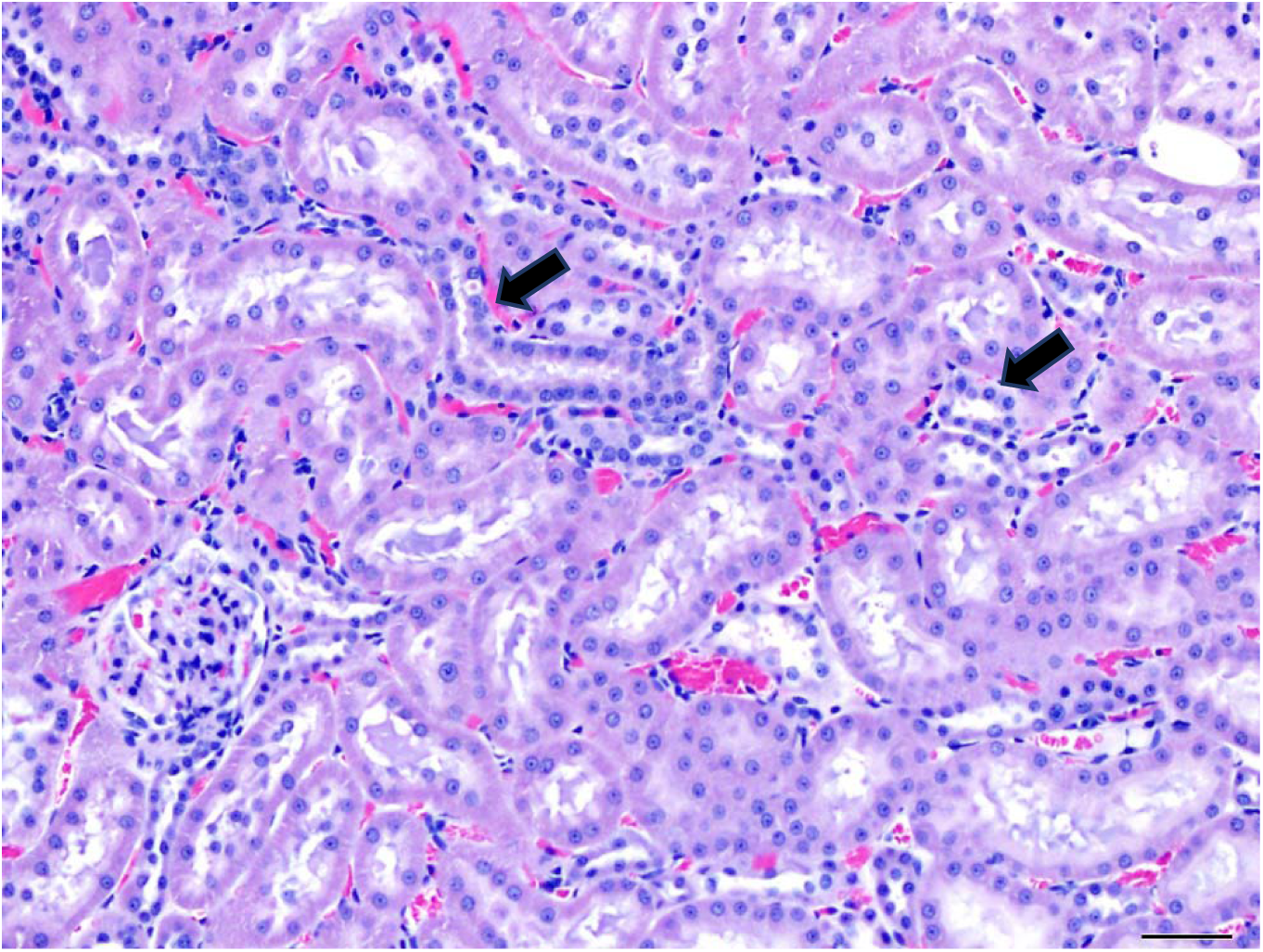
Control group: renal tubular regeneration (arrows); score = 2.

